# Nutrient currencies and P resorption greatly amplify the perceived costs of reproductive allocation in perennial heath shrubs

**DOI:** 10.64898/2026.06.10.731282

**Authors:** Elizabeth Wenk, Daniel Falster, Ian Wright, Mark Westoby

## Abstract

**Summary:**

- In woody perennials, reproductive allocation considered as a fraction of NPP (RA) has been rarely quantified, especially tracked across plant lifetimes, measured in biologically meaningful currencies, and separated from standing biomass.
- We addressed this gap by measuring dry mass, nitrogen, and phosphorus for every aboveground tissue type for 14 iteroparous perennial shrubs tracked across their full lifetimes, calculating RA using four accounting schemes and three currencies.
- RA was substantially higher by mid-life than typical estimates from ecosystem-scale studies. Distinguishing standing biomass from yearly production further revised RA upwards. Using a nutrient currency (especially P) increased the perceived costs of reproduction. Propagules were highly nutrient-enriched and reproductive accessory costs consumed more nutrients than did the propagules themselves. Considering N and P resorption from senescing leaves and wood shrank the effective vegetative nutrient budget, further concentrating net annual nutrient demand in reproductive tissues.
- Our results highlight high investment in RA for woody perennials, especially using nutrient currencies. Broadly similar allocation patterns were observed across species with different functional traits and lifespans, suggesting generality that may apply across biomes. Widespread underestimation of RA in forest growth models likely overestimates the proportion of NPP available for vegetative growth, leading to substantial errors in predictions.

## Introduction

How much energy plants allocate to reproduction versus growth remains a central unresolved question in life history theory and ecosystem ecology. Carbon fixed through photosynthesis must be partitioned among tissues that differ in biochemical composition, turnover rates, and ecological roles, with consequences for food web dynamics and nutrient cycling. Allocation to reproduction inevitably trades off against vegetative growth and species differ in how this trade-off translates into different life histories (Stearns, 1992; Obeso, 2002; Wenk & Falster, 2015; Wenk *et al*., 2018; Dorken *et al*., 2025; Ward *et al*., 2025). Further, the magnitude of investment could depend on what currency measures investment (Reekie & Bazzaz, 1987; Ashman & Baker, 1992; Pérez-Martínez & Méndez, 2021). Reproductive costs are most-commonly expressed in units of biomass, but seeds and reproductive tissues are disproportionately enriched in nitrogen and phosphorus relative to vegetative organs — suggesting that nutrient-based currencies may reveal substantially higher reproductive costs than biomass alone implies (Reekie & Bazzaz, 1987; Obeso, 2002; Pérez-Martínez & Méndez, 2021).

To date, research on reproductive allocation (RA) has been dominated by herbaceous plants (Obeso, 2002; Wenk & Falster, 2015; Pérez-Martínez & Méndez, 2021), particularly focussing on distinction of reproductive strategies and effects of size on reproductive output (Klinkhamer *et al*., 1992; Weiner *et al*., 2009; Bonser & Aarssen, 2009). Herbs are easy to study, with annuals completing their life cycle in a single year and many perennials retreating to underground storage organs during an unfavourable growing season. Annuals have the advantage that they can easily be grown under experimental conditions, are small enough to easily harvest and separate into tissue types, and often display distinct shifts from growth to reproduction as they age (Reekie & Bazzaz, 2005). By contrast, woody perennials may live tens to thousands of years, gradually shift their relative allocation to growth versus reproduction across time, and have aboveground vegetative tissues that have formed across many years. Yet herbaceous species are only a small part of global biodiversity and ecosystem energy flows; seventy percent of the world’s standing plant biomass is wood (Erb *et al*., 2018; Bar-On *et al*., 2018), and yet more if the non-wood tissues of woody perennials are included.

Our understanding of RA in woody perennials rests largely on ecosystem-scale biomass accounting, where RA can be inferred from litterfall — and by this measure appears modest, averaging less than 0.2 of NPP across biomes (Malhi *et al*., 2011; Kitayama *et al*., 2015; Hanbury-Brown *et al*., 2022; Ward *et al*., 2025). Many studies measure RA by counting litterfall for an entire forest, rather than for individual trees (Bullock, 1995; Malhi *et al*., 2011; Kitayama *et al*., 2015; Wright *et al*., 2024; Gougherty & Templer, 2024). However, these accounts average across life stages and species and do not separate genuinely new growth from turnover, meaning it is difficult to get a fitness-focussed estimate of RA. By contrast, few studies track individual level estimates of RA in perennial plants (but see Delerue *et al*., 2013; Edwards *et al*., 2015 for recent efforts). For fourteen heath shrub species studied across their full lifetimes, we previously showed that reproductive allocation increases with plant age and that nearly all species approach an RA of 100% of surplus energy — that is, energy that remains after accounting for replacement of pre-existing tissues — as growth slows in older individuals (Wenk *et al*., 2018; Dun *et al*., 2025). These findings suggest that at the right life stage, woody perennials invest substantially more of surplus energy to reproduction than total ecosystem accounts indicate (Pérez-Martínez & Méndez, 2021; Ward *et al*., 2025), and much closer to the RA of annuals and many other organisms (Hirshfield & Tinkle, 1975; Rocha *et al*., 2001), as well as expectations of eco-evolutionary theory (Iwasa & Cohen, 1989; Kozlowski, 1992; Wenk & Falster, 2015).

Whether this high reproductive investment increases or decreases when RA is measured in nutrient rather than biomass currencies — and whether conclusions are stable across different reproductive accounting schemes — remains unresolved (Reekie & Bazzaz, 1987; Ashman, 1994; Kerkhoff *et al*., 2006; Pérez-Martínez & Méndez, 2021). Plant tissues differ systematically in nutrient concentration based on their function: wood is structural and nutrient-poor, leaves are metabolically active and nitrogen-rich, and propagules carry the highest concentrations of all, especially phosphorus, to provision young seedlings (Obeso, 2002; Elser *et al*., 2010). Reproductive accessory tissues show equally diverse forms to vegetative tissues, yet their nutrient concentrations are rarely measured separately; the few studies that have done so find strong differences among tissue types (Groves *et al*., 1986; Groom & Lamont, 2011; Edwards *et al*., 2015). Nutrient resorption from senescing leaves further complicates accounting, reducing the effective vegetative nutrient cost relative to reproductive tissues (Killingbeck, 1996; Edwards *et al*., 2015). Together, these factors suggest that shifting to a nutrient currency would likely increase estimated reproductive allocation — but the direction and magnitude of that shift for woody perennials remains sparingly quantified (Reekie & Bazzaz, 1987; Ashman & Baker, 1992; Pérez-Martínez & Méndez, 2021). Estimates of RA can also depend meaningfully on the method of accounting (Box 1). Whether investment is measured from a single harvest or tracked across a lifetime and whether standing biomass or annual production is summed into the denominator can shift conclusions substantially, particularly for iteroparous perennials where reproductive effort increases with age and tissues can persist for many years. Thus different methodological approaches can seemingly impact how we interpret empirical data and apply results to understand the evolution of plant life histories.

Here we address these uncertainties by measuring dry mass and nutrient investment across all aboveground tissue types for fourteen perennial heath shrubs, tracked across their full lifetimes. We ask two questions: how do nitrogen and phosphorus concentrations vary among tissue types, including reproductive accessory tissues and senesced vegetative tissues; and does shifting from a biomass to a nutrient currency — or among the several accounting schemes described in Box 1 — change the conclusions about reproductive allocation established by prior work in this system? Figure 1 summarises how these factors combine: relative to a standard single-harvest biomass estimate, accounting for plant age, shifting to nutrient currencies, and incorporating resorption each increase the perceived contribution of reproductive tissues — suggesting that current biomass-based estimates of RA in woody perennials are likely to be conservative.

**Figure 1.**
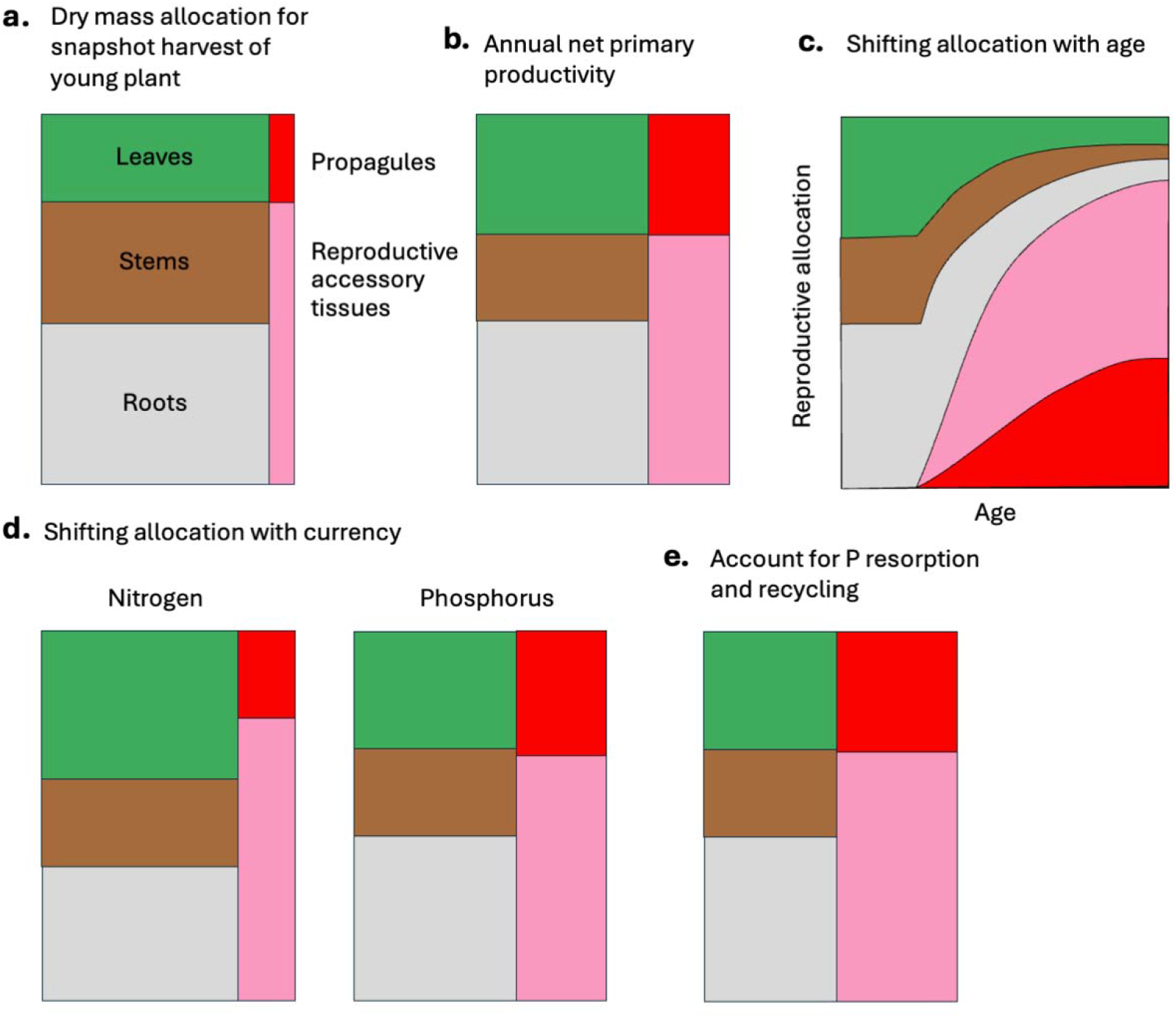
Schematic showing how plant age, allocation currency, and allocation calculations can impact perceived relative allocation to vegetative and reproductive tissues. a) Studies most commonly report a single biomass harvest, with relative allocation measured as the standing biomass of each tissue type at the time of harvest; b) Separating standing biomass from annual growth reduces the relative allocation to woody tissues and leaves, especially leaves with long leaf life spans; c) Older plants will have a higher proportion of reproductive biomass than younger plants; d) Calculations using nitrogen or phosphorus concentrations will indicate a relatively lower investment in vegetative materials relative to reproductive tissues; and e) Accounting for nutrient resorption will further reduce the relative contribution of vegetative tissues, as plants can resorb nutrients more effectively from these than most reproductive tissues. See Box 1 for further details.

#### Box 1. Description of alternative accounting schemes

Different accounting schemes can shift the perceived relative investment into different tissues. Resource allocation, whether measured using a dry mass or nutrient currency, is most commonly assessed from a single “snapshot” harvest, with standing biomass divided into different tissue pools (Figure 1a) (Weiner, 2004; Wenk & Falster, 2015; Pérez-Martínez & Méndez, 2021). Factors including allocation currency, plant age, accounting for biomass formed during a previous growing season, and accounting for resorption will all change the relative magnitude of each tissue pool. For instance, a single harvest cannot separate yearly growth (NPP) from pre-existing standing biomass, and subtracting the vegetative tissues formed during a previous growing season will increase RA (Figure 1b). For perennial, iteroparous species, those with multiple reproductive episodes across their lifetime, reproductive allocation will generally increase with plant size and age (Figure 1c) (Weiner, 2004; Wenk *et al*., 2018; Ward *et al*., 2025). Changing to nutrient currencies will likely indicate higher allocation to reproductive tissues, especially if calculations use P concentrations (Figure 1d) (Reekie & Bazzaz, 1987; Obeso, 2002; Pérez-Martínez & Méndez, 2021). For many species, nutrient resorption, especially from senescing leaves, offers a key input to their total nutrient budget (Killingbeck, 1996; Edwards *et al*., 2015; Dhakal *et al*., 2025), but calculating the magnitude of this pool requires measurement of nutrients and dry mass in live and senesced tissues and knowledge of leaf lifespan or the mass of leaves shed each year (Figure 1e) (Wright & Westoby, 2003).

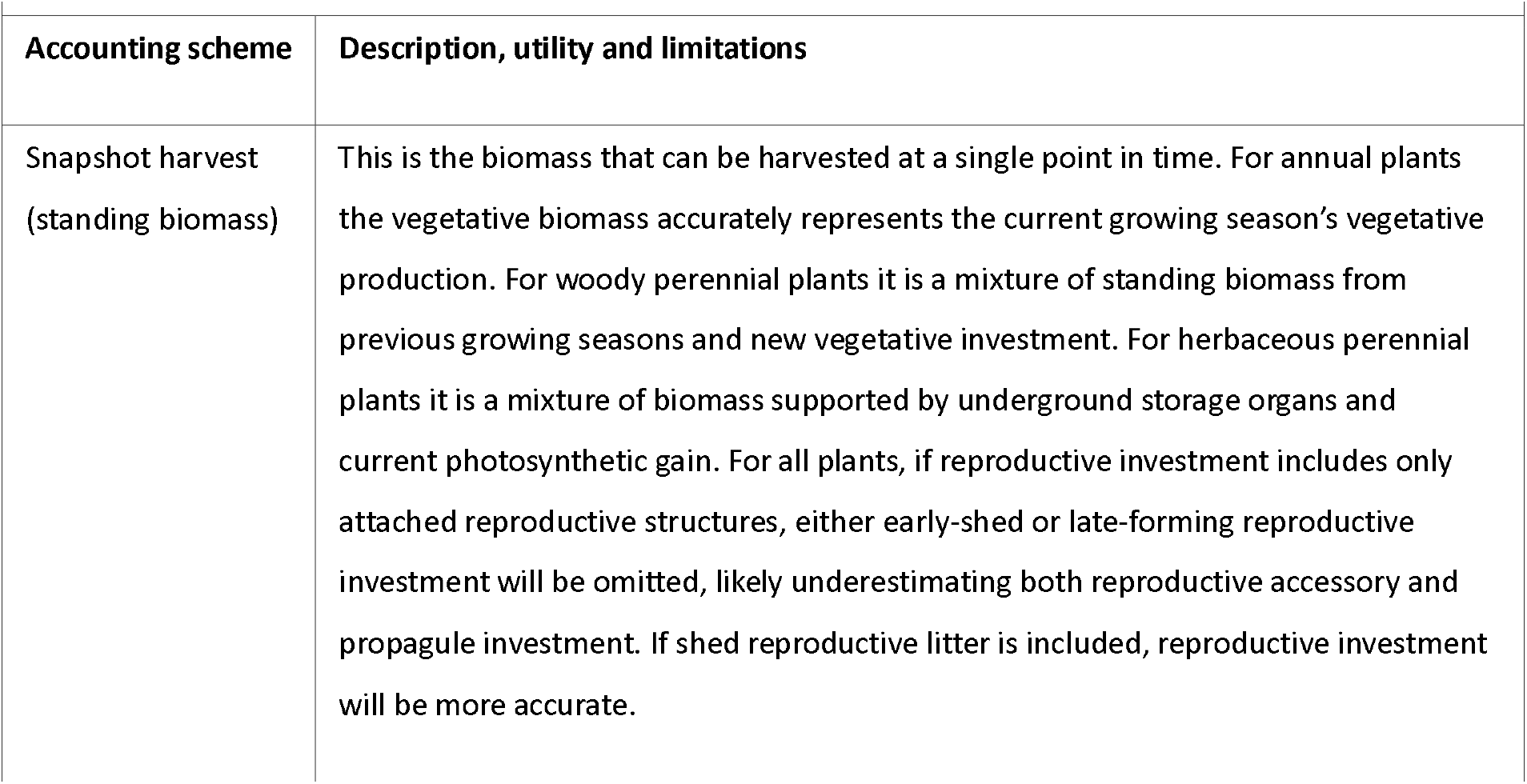
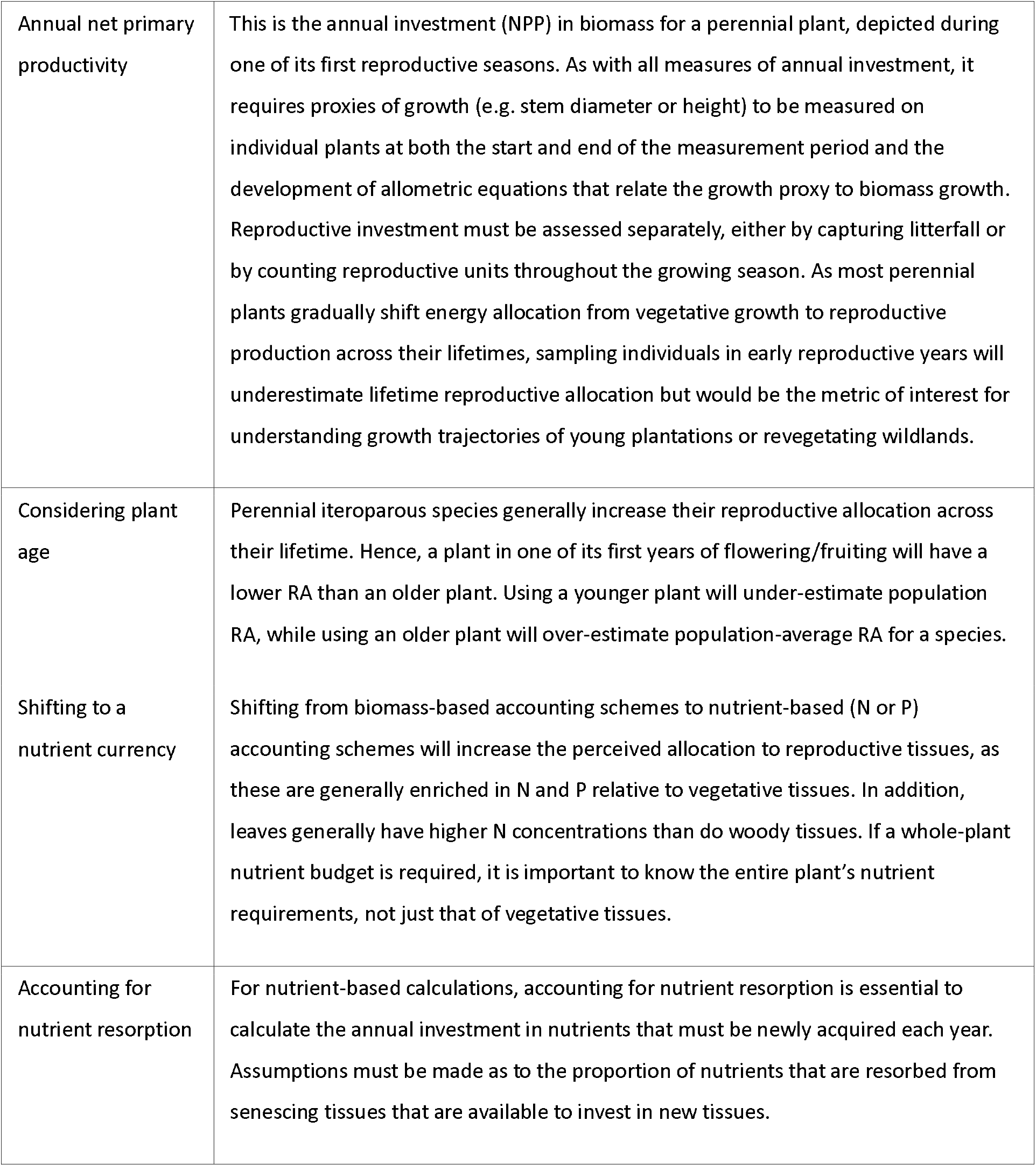

## Materials and Methods

### Study system

We conducted our research on 14 common woody, perennial, monoecious, iteroparous shrub species in a fire-prone coastal heathland in Kuring’gai Chase National Park, northeast of Sydney, Australia (Kodela & Dodson, 1988). All species selected are obligate seeders, killed by hot fires and reestablishing from seed, generally within a year of a fire. As such, the age of individual plants could be accurately determined from the time of the most recent fire, based on fire records maintained by the New South Wales National Parks and Wildlife Service and using time-since-fire as a proxy for plant age. Patches of vegetation were selected at six different times since fire (harvested at 1.35 years, 2.4 years, 5 years, 7 years, 9 years, and 32 years; Table S1). Within the mapped burn areas, study sites were chosen at locations where no individual plants of the study species were notably taller than other individuals, indicating the fire intensity was sufficiently high to burn all standing vegetation and ensuring only post-fire germinated individuals were selected. At the selected sites, *Banksia ericifolia, Hakea teretifolia* and *Allocasuarina distyla* (not included in our study because it is dioecious) would be the dominant canopy species late in succession, at heights of 3–5 m (Carolin & Tindale, 1993). Nearby locations included a *Eucalyptus* canopy, but sites were chosen to have minimal *Eucalyptus* cover, and all study individuals were chosen to be unshaded by *Eucalyptus* across the middle of the day, as these large post-fire resprouting trees tend to shade out the long-lived mid-canopy obligate reseeders, notably changing the species composition from adjacent landscape patches lacking *Eucalyptus* cover.

To measure all aboveground biomass investment, we used a sampling design that allowed us to tabulate gross and net investment in all aboveground tissue types across a year of growth. Seven healthy individuals of each species were selected at each site for sites aged 16 months or greater and 14 seedlings at the youngest site, as seedlings experience a higher mortality rate. Plants were selected during May–June 2012, months of minimal growth and reproductive investment in the plant community and followed for one year. One additional recently burned site was monitored from June 2013 to June 2014 and the youngest of the previous sites was monitored for a second year with a new cohort of plants, also from June 2013 to June 2014. A total of 597 tagged individuals were alive at the end of the study period and form the basis for the data collected.

### Biomass measurements

Vegetative investment into leaves and stems was determined across the one-year study period. At both the start and conclusion of the study, we recorded the height and basal diameter (measured at 10 cm above ground-level in plants aged 3+ and at 1 cm in seedlings) of each individual. All plants were harvested at the conclusion of the study, then separated into stems, leaves, and reproductive tissues. Roots could not be harvested for these species. Plant parts were oven dried at 60ºC for at least one week before dry mass was determined. Log-log allometric lines were fitted to the stem diameter-stem mass and plant age-leaf mass data for each species. For each individual, yearly net increase in stem and leaf mass was calculated from these curves, based on the individual’s stem diameter and age increments (described fully in Wenk *et al*., 2018). In addition, leaf turnover was calculated on a selected leading shoot, with leaf number counted at the start of the study period and the number of remaining leaves counted at the study’s conclusion (Wright *et al*., 2002). For species with small leaves that attached to the stem in a regular pattern, leaf count was determined from measurements of “stem length covered by leaves” for the selected shoot and “leaves per cm stem length” for the species. The calculated leaf turnover value was applied to the entire plant.

To assess reproductive investment, plants were visited every four weeks during winter and autumn and every three weeks during spring and summer. At each visit, all flowering and fruiting materials were counted on each individual, including buds (by size class), flowers, immature fruit (by size and maturity classes), mature fruit and other reproductive structures specific to certain species, such as woody cones. For some species, the dimensions of immature and mature fruit and cones were also measured, as the final size of the structures was quite variable. The exact flowering parts included varied considerably by species, due to their diverse floral structures (see Supplementary Information in Wenk *et al*. 2018 for figures indicating reproductive structures censused for each species). The census timing ensured that individual reproductive parts did not progress through more than one reproductive stage (i.e. bud, flower, immature fruit, mature fruit) between visits and the progression of each reproductive unit could be followed until abortion or maturation. Reproductive parts on nearby untagged plants were simultaneously collected to determine biomass of each reproductive part. Total reproductive biomass on an individual was determined by summing the count of each reproductive unit at the most developed stage it achieved by that part’s dry biomass; that is, the count of aborted buds was multiplied by bud mass, the count of flowers (aborted and those with ovaries that developed into fruit) by flower mass, and the count of mature fruit by fruit mass.

### Tissue nutrient concentrations

Nitrogen and phosphorus concentrations were determined on green leaves, wood, bark, senesced leaves, shed wood (senesced twigs), and shed bark from 2 – 4 of the harvested individuals (randomly selected) of each species at each site. Two types of wood were analysed: twigs, representing sapwood, and basal stems, representing a mixture of sapwood and heartwood. Nutrient concentrations were also measured on a selection of reproductive tissues for each species, with 2 – 4 replicates collected across sites but bulked for analysis. Nutrient concentrations were independently measured for flowers (including buds), green floral parts (bracts, calyces), immature fruits, mature fruits/seeds/propagules (depending on species), and woody cones, with additional subdivisions of tissues for some taxa. For some reproductive tissues and for senesced leaves for some species, samples were bulked from many individuals to achieve a sufficiently large sample size. Tissue nitrogen concentration was determined by combustion using a LECO TruSpec CHN analyser. For phosphorus, tissue samples were digested in acid and total P concentration determined by inductively coupled plasma optical emission spectrometry (ICP-OES).

For each plant, for each tissue type, annual nitrogen and phosphorus investment was calculated as the product of nutrient concentration and the corresponding tissue dry mass investment. Leaf nutrient investment was calculated both including and excluding resorption: 1) for calculations ignoring resorption, yearly gross leaf investment was multiplied by green leaf nutrient concentration; 2) for calculations accounting for resorption, yearly leaf nutrient investment was calculated as the sum of the product of net leaf investment by green leaf nutrient concentrations and the product of shed leaf mass by nutrient loss from senesced tissues (green leaf nutrient concentration – senesced leaf nutrient concentration). Although nutrient concentrations of both live and shed sapwood and bark were measured, only live nutrient concentration was considered for these tissues due to our inability to reliably separate stems into sapwood and heartwood components.

### Allocation calculations

Relative allocation to four tissue categories was calculated using four different accounting schemes and three currencies (dry mass, N concentration, P concentration). The four tissue categories used were: leaves, stems (including sapwood, heartwood, and bark), propagules (generally seeds, but entire fruit for species where the seed was fused within a small, dry fruit), and accessory reproductive tissues. Accounting schemes were chosen to reflect those most commonly used in the literature and those best suited to address different research questions:

1. **Snapshot harvest**. This is the standing biomass at a single point in time, calculated on a mid-aged plant and is the method most commonly reported in the literature, as only a single visit to a site is required. It is the appropriate accounting scheme for research questions focused on current site biomass. Our calculations used the year-end standing vegetative aboveground biomass and total reproductive investment across the year.
2. **Annual growth investment in a young reproductively mature plant**. Our calculations used the wood and leaf investments derived from our allometric equations and total reproductive investment across the year. For species with lifespans of ~10 years, data from the 2.4-year-old site were used and for species with lifespans ~20+ years, data from the 6-year-old site were used (site age referring to time since last fire).
3. **Annual growth investment in a mid-aged plant**. Our calculations used the wood and leaf investments derived from our allometric equations and total reproductive investment across the year. For species with lifespans of ~10 years, data from the 6-year-old site were used and for species with lifespans ~20+ years, data from the 8-year-old site were used.
4. **Annual growth investment in a mid-aged plant, accounting for nutrient resorption from shed leaves**. Our calculations were identical to those above, but the nutrients required to produce leaves were those that will eventually be shed with the newly produced leaves. Nutrients that are resorbed from senescing tissues are assumed to be 100% available to invest in new tissues. Resorption from bark and stems is ignored, as we were unable to accurately predict nutrient resorption as sapwood transitions to heartwood. Resorption from reproductive tissues is unknown, but it would only apply to floral reproductive parts that undergo planned senescence.

To capture how the cost of tissues shifts with currency, for each accounting scheme and each species, we calculated the ratio of the tissue cost in each nutrient currency to the tissue cost calculated using dry mass. Tissues with a relative cost > 1 become relatively costlier to the species when shifting to a specific nutrient currency, while values < 1 become relatively less costly.

### Statistical analyses

To assess whether live vegetative tissues differed in nutrient concentration, linear models were fit with species and tissue type as predictor variables and log nutrient (N and P) concentration as the response variable. These models could only include vegetative tissues, as nutrient concentration was assessed on a single bulked tissue sample for each reproductive tissue within each species. To test for differences in nutrient concentration across reproductive tissues a linear mixed-effects model was fit using the lme4 R package (Bates *et al*., 2003), with accessory tissue type as a fixed effect and species as a random effect.

To determine the N to P scaling relationship across species within each vegetative and reproductive tissues category (leaves, wood, bark, green accessory tissues, floral accessory tissues, woody accessory tissues, and propagules), a standardised major axis regression was fit to N and P concentrations for each tissue type for which the two variables were correlated, using the smatr R package (Warton *et al*., 2006).

To compare allocation across tissue categories and across accounting schemes, the proportional allocation to each vegetative and reproductive tissue category was calculated using biomass, nitrogen and phosphorus currencies for each of the four accounting schemes. All analyses were completed using R v4.4.0 (R Core Team, 2024).

## Results

Two streams of information must be integrated to calculate RA for each plant and under different currencies and accounting scheme: 1) nutrient concentrations of live and senesced tissues; and 2) total annual investment into each tissue type.

### Tissue nutrient concentrations

Overall, we found that tissue level nutrient concentrations varied considerably between species for wood, bark, propagules and accessory tissues (less so among leaves); also, that the average concentrations varied much more between tissue types than they did between species (Table 1). Nitrogen and phosphorus concentrations in different tissues and species of this heath community are summarized in Figure 2a.

**Table 1.** Nitrogen and phosphorus concentrations of live tissues, across all species. Reproductive accessory tissue nutrient concentrations are weighted by the relative concentration of all tissues to each species total investment. Values in brackets are ranges across species; see Table S1 for values for each species.

| Live Tissue | N<br>(mg/g ± SE) | P<br>(mg/kg ± SE) | N:P |
| --- | --- | --- | --- |
| Leaves | 12.7 ± 0.86<sup>d</sup><br>(7.40—19.36) | 213 ± 15.7<sup>c</sup><br>(129—391) | 63.6 ± 4.0<sup>b</sup><br>(41.0—87.9) |
| Sapwood | 2.96 ± 0.36<sup>b</sup><br>(1.36—5.99) | 110 ± 10.4<sup>d</sup><br>(70.6—218) | 32.8 ± 3.9<sup>a</sup><br>(13.0—60.1) |
| Wood from base | 2.14 ± 0.36<sup>a</sup><br>(0.234—4.22) | 67.3 ± 11.2<sup>e</sup><br>(37.4—207) | 49.5 ± 7.5<sup>ab</sup><br>(4.42—90.6) |
| Bark | 5.69 ± 0.67<sup>c</sup><br>(2.55—11.0) | 118 ± 6.9<sup>d</sup><br>(85.7—172) | 60.4 ± 7.9<sup>b</sup><br>(22.2—110) |
| Reproductive accessory tissues | 12.1 ± 0.94<sup>d</sup><br>(5.56—17.1) | 502 ± 52.6<sup>b</sup><br>(182—775) | 37.4 ± 45.2<sup>a</sup><br>(12.7—79.2) |
| Propagules | 44.1 ± 6.41<sup>e</sup><br>(5.55—96.1) | 4253 ± 1464<sup>a</sup><br>(93.3—21455) | 38.3 ± 16.7<sup>ab</sup><br>(3.64—201) |

**Figure 2.**
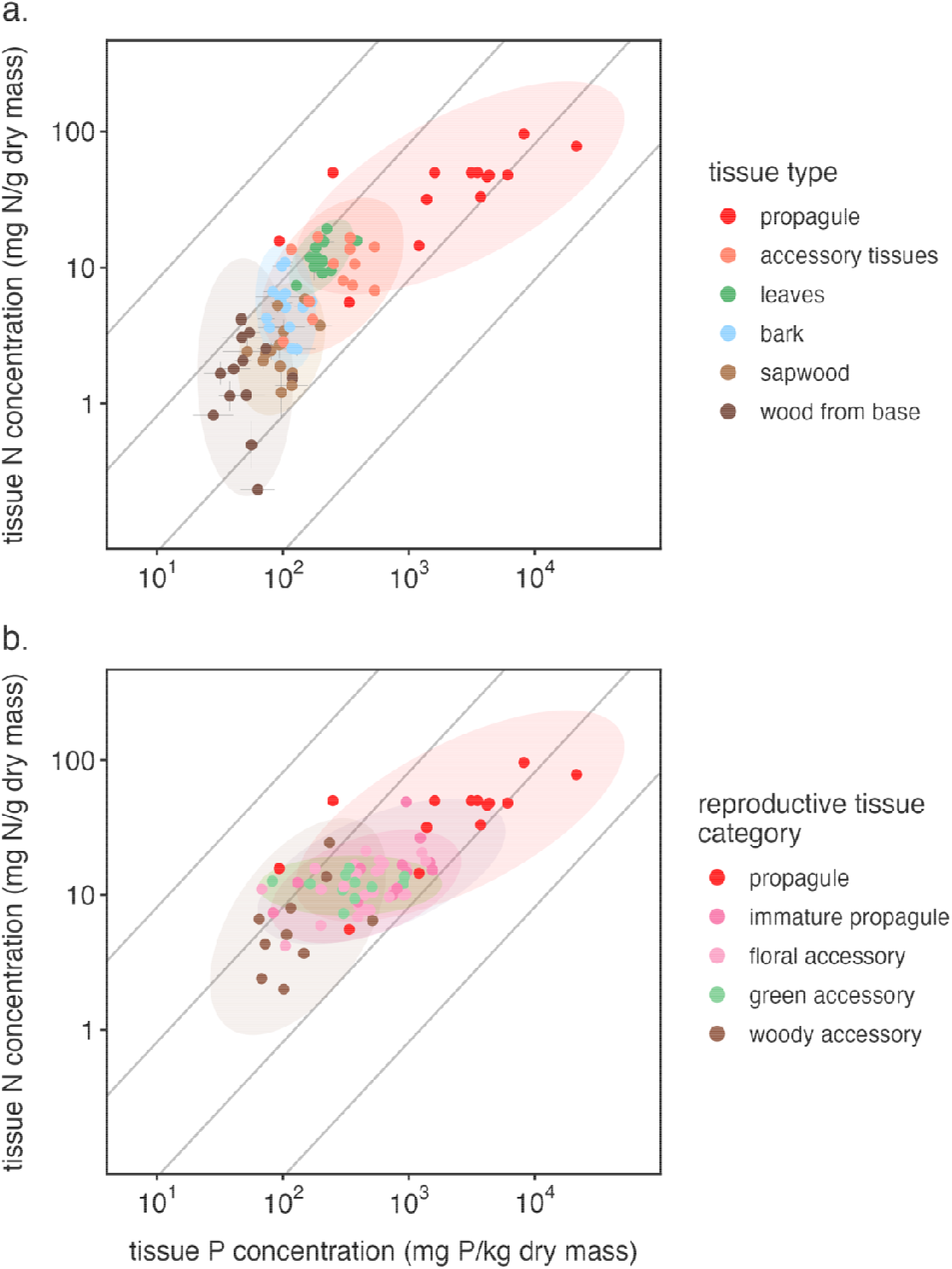
Tissue nutrient concentrations, indicating that although there is variation across species, the between tissue variation is much greater. Grey lines indicate 1:1 N:P scaling. Panel (a) compares different reproductive and vegetative tissue types, with all accessory tissues merged into a single “accessory tissues” values, while panel (b) separates the different categories of accessory tissues. The 95% confidence ellipses around expected species mean N and P concentrations by tissue type are shaded.

First, tissue types differed strongly in nitrogen concentration (F_5,350_ = 394, p < 0.001), with much smaller, but still highly significant effects of species (F_13, 350_ = 14.9, p < 0.001), and a tissue type by species interaction (F_52, 350_ = 5.12, p < 0.001) (Tables 1, S2, Figure 2a). Further, all pairwise comparisons of nitrogen concentration between tissue types were highly significantly different (p < 0.001), except leaves versus accessory tissues (p = 0.088). Phosphorus concentration was likewise highly significantly different across tissue types (F_5,350_ = 181, p < 0.001), with smaller, but still significant effects of species (F_13, 350_ = 2.66, p = 0.0014), and a tissue type by species interaction (F_52,350_ = 2.56, p < 0.001) (Tables 1, S1, Figure 2a). All pairwise comparisons between tissue types were highly significantly different (p < 0.001), except sapwood versus bark (p = 0.362).

Second, N:P ratio varied widely among tissues (Tables 1, S2), with means ranging from 32.8 to 63.6 among tissue types (Table 1), but more broadly from 13.1 to 109.8 across species by tissue type (Table S2). These differences were highly significant for tissue (F_5,350_ = 14.13, p < 0.001) and species (F_5,350_ = 5.21, p < 0.001), with a slightly weaker tissue type by species interaction (F_5,350_ = 1.70, p = 0.0013). The N:P ratio was significantly higher in the more metabolically active tissues, leaves and bark, compared to sapwood and reproductive accessory tissues. N and P concentrations across species were uncorrelated in sapwood, wood from the base of the stem, bark, and reproductive accessory tissues (Fig 2a), marginally correlated in leaves (p=0.058, R^2^=0.27), and strongly correlated in propagules (p=0.006, R^2^=0.47). The study species’ leaf N:P scaling relationship (Figure 2a) was indistinguishable from the N ∝ P^2/3^ relationship (mean = 0.94, 95% CI = [0.56, 1.58]). Propagules displayed a steeper scaling relationship, with P concentration increasing more rapidly than N in the propagules of the study species (mean = 1.94, 95% CI = [1.27, 3.07]).

Whereas reproductive accessory tissues as a whole show an intermediate nutrient content (Fig 2a), further subdividing these into different categories reveals greater diversity of nutrient concentrations among tissue types (Fig 2b). The individual tissues that contribute to total reproductive accessory biomass spanned broad ranges of N and P concentrations (Tables 2, S3, Figure 2b). These tissues were divided into four categories, woody reproductive tissues (cones, woody seed pods), green reproductive tissues (sepals, calyx, other green tissue), floral reproductive tissues (buds, flowers), and immature fruits (developing, but not yet mature). These tissue types differed significantly in both N concentration (F_4, 16_ = 18.8, p < 0.001) and P concentration (F_4, 16_ = 17.8, p < 0.001), but for both models the species effects (F_13, 16_ = 1.80, p = 0.13 for N and F_4, 16_ = 1.30, p = 0.30 for P) and interactions (F_40, 16_ = 0.71, p =0.81 for N and F_4, 16_ = 1.43, p = 0.22 for P) were non-significant and were dropped from the model before pairwise tissue comparisons were performed. Pairwise comparisons indicated that propagules had significantly higher N and P concentrations than any accessory tissue type, that woody accessory tissues had significantly lower N and P concentrations than any other accessory tissue type, and that immature propagules, floral accessory tissue, and green accessory tissues had N and P concentrations that were statistically indistinguishable from each other (Table 2). Despite propagules having dramatically higher N and P concentrations than other accessory tissues, their N:P ratio was not distinct (F_4, 69_ = 0.6238, p = 0.6471). While N and P concentrations were uncorrelated across study species for woody accessory tissues, they were significantly correlated for both immature propagules and floral accessory tissue; both had N:P scaling slopes steeper than N ∝ P^2/3^ (immature propagules: mean = 1.83, 95% CI = [1.05, 3.20]; floral accessory tissues: mean = 1.80, 95% CI = [1.25, 2.59]).

**Table 2.** Nutrient concentrations of specific reproductive tissues, across all species. The values for propagules are identical to those in Table 1. Values in brackets are ranges across species; see Table S2 for values for each species.

| Reproductive Tissue Type | N<br>(mg/g) | P<br>(mg/kg) | N:P |
| --- | --- | --- | --- |
| Propagules | 44.1 ± 6.41<sup>a</sup><br>(5.55—96.1) | 4253 ± 1464<sup>a</sup><br>(93.3—21455) | 38.3 ± 16.7<br>(3.64—201) |
| Immature propagules | 17.1 ± 3.2<sup>b</sup><br>(7.38—49.1) | 802 ± 139<sup>b</sup><br>(83.5—1546) | 33.3 ± 8.6<br>(9.86—94.7) |
| Floral accessory tissues | 12.4 ± 0.9<sup>b</sup><br>(6.90—19.2) | 556 ± 63<sup>b</sup><br>(276—904) | 32.5 ± 6.0<br>(12.3—72.7) |
| Green accessory tissues | 12.0 ± 0.7<sup>b</sup><br>(7.29—14.9) | 416 ± 81<sup>b</sup><br>(164—926) | 44.5 ± 12.1<br>(13.3—88.3) |
| Woody accessory tissues | 7.7 ± 2.1<sup>c</sup><br>(2.40—13.6) | 164 ± 43<sup>c</sup><br>(64.7—515) | 53.8 ± 10.2<br>(12.6—102) |

### Nutrient resorption

High proportions of P were resorbed from leaves, sapwood, and bark before senescence (Table 3, Figure 3), while far lower proportions of N were resorbed. The degree of P resorption was relatively similar across species, ranging from 56.9% to 97.9% for sapwood, 12.6% to 73.2% for bark and 59.3% to 92.0% for leaves. By contrast, N resorption spanned from 0% to 60.9% for sapwood, 0% to 29.1% for bark and 0% to 76.7% for leaves. For many species, N concentration was higher in senesced leaves than in live leaves (Table S4), such that across the community NUE was very low.

**Table 3.** Nutrient resorption from vegetative tissues, across all species. See Table S3 for values for each species.

| Senesced Tissue | N<br>(mg/g ± SE) | N<sub>resorb</sub><br>(%) | P<br>(mg/kg ± SE) | P<sub>resorb</sub><br>(%) | N:P |
| --- | --- | --- | --- | --- | --- |
| Leaves | 0.961 ± 0.070 | 21.0 | 52.1 ± 7.64 | 76.6 | 274 ± 34.9 |
| Sapwood | 0.249 ± 0.019 | 5.77 | 19.0 ± 2.04 | 79.2 | 32.8 ± 40.8 |
| Bark | 0.569 ± 0.048 | 1.19 | 56.1 ± 3.73 | 49.4 | 60.4 ± 14.9 |

**Figure 3.**
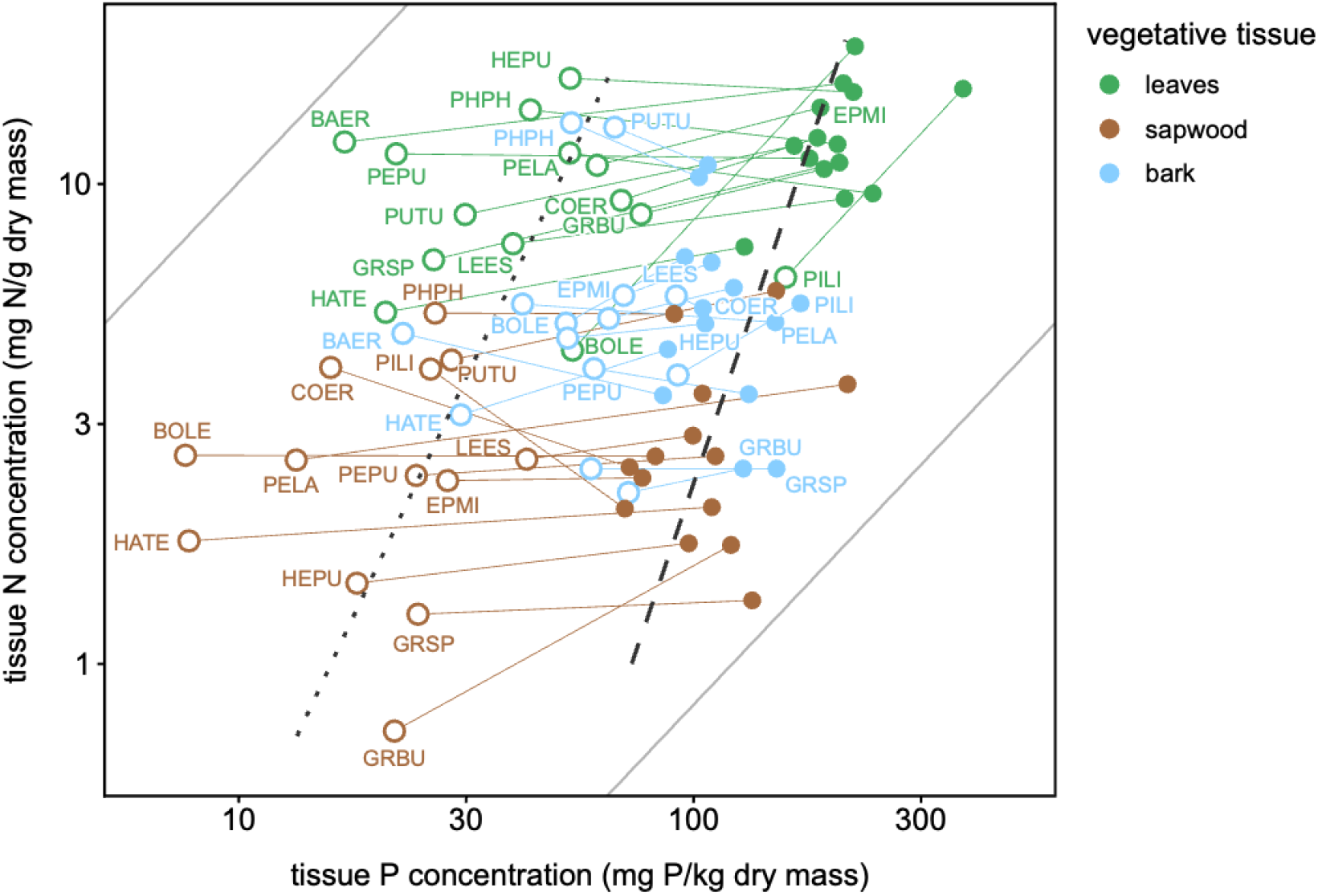
Nutrient resorption from vegetative tissues, with closed symbols indicating nutrient concentrations of live tissues and open symbols nutrient concentrations of senesced tissues. Grey lines indicate 1:1 N:P scaling. The dashed black line is N:P scaling across live tissues and dotted black line is N:P scaling across senesced tissues.

### Tissue allocation calculations

Allocation patterns looked very substantially different depending on accounting scheme (top to bottom in Figure 4) and on resource currency (left to right in the bar graphs of Fig 4), with quite modest differences between species (Figure 4, Table S5). The righthand column indicates the shift in proportional costs, relative to dry mass. That is, points above 1 represent tissues that become more costly when calculating using a specific nutrient currency, while those below 1 are tissues that become less costly using nutrient currencies.

**Figure 4.**
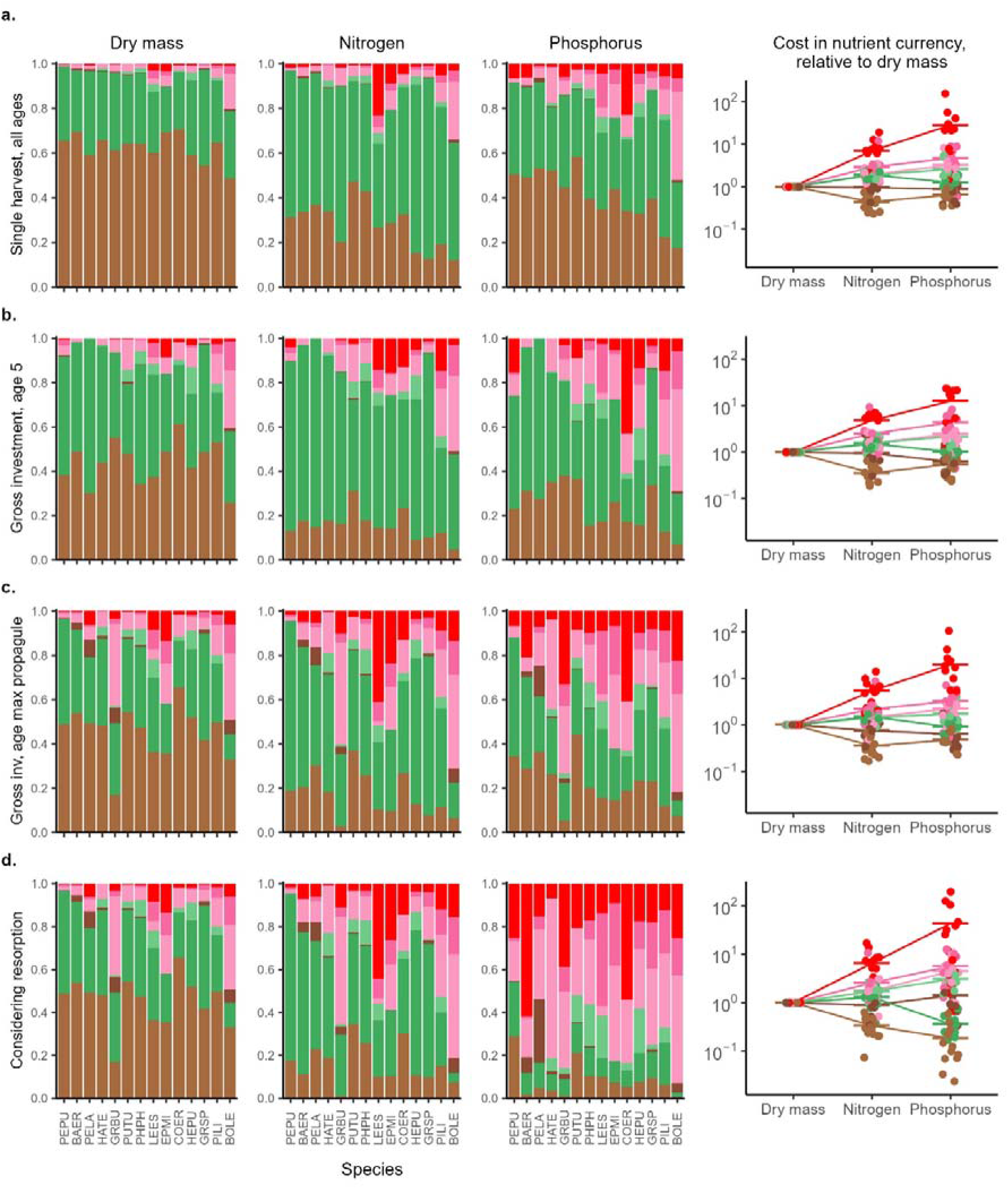
Distribution of investment in propagules, specific accessory tissues (aborted immature propagules, floral tissues, green tissues, and woody tissues), leaves and sapwood (including bark) across species, across currencies, and using different investment metrics. Species are arranged based on a combination of lifespan and their maximum relative investment in reproduction across all metrics. Row (a) are calculations based on a snapshot harvest, row (b) are plants in their first year(s) of reproducing, row (c) are plants at an age with high reproductive allocation, and row (d) incorporates nutrient resorption from leaves. The first column displays allocation calculations on a dry mass basis, the second column using nitrogen concentrations, the third column using phosphorus concentrations, and the fourth column indicates each tissue’s cost in nutrient currencies relative to its cost measured using dry mass. Each point represents a species’ mean for a given tissue, jittered in the x-dimension for visibility.

Standing harvest calculations (Figure 4a) included many years of wood growth and generally several years of leaf production, with only a single year of reproductive investment. Consequently, RA appears low, ranging from 1.2% to 21.3%.

In biomass, older plants (Figure 4b) allocated a greater proportion of their biomass investment to reproduction than did younger plants (Figure 4c). There was relatively little variation in the proportional allocation across species, with the exception of the shortest-lived species on the far right of the bar graphs (*Pimelea linifolia* and *Boronia ledifolia*) which both had a high ratio of reproductive to vegetative investment from a young age.

Shifting to N currency (i.e., comparing the first to the second column of panels in Figure 4) accentuated the allocation to leaves relative to wood, due to their higher N concentration (Figure 2a) and also increased the relative allocation to all accessory tissues and propagules.

Shifting to P currency (third column, Figure 4) implied a much higher investment in both propagules and accessory tissues, across all species, reflecting the elevated P content of reproductive tissues relative to vegetative tissues.

Accounting for nutrient resorption, further accentuated the relative allocation to reproductive versus vegetative tissues. As leaves (and wood) resorb high proportions of P before senescence, this shift in allocation is especially apparent in Figure 4d.

In the plots of P investment (third column), the high cost of floral tissues and aborted fruit was accentuated; for most species, these investment pools were much larger than nutrient investment into propagules, even at the age of maximum propagule output, due to their high biomass contribution. Across nearly all species, propagules and reproductive accessory tissues become more costly, while woody tissues are cheaper, with stronger shifts for P (RA from 45.6—9.3%) than for N (RA from 4.6—88.2%). Leaves and green reproductive tissues show more muted and variable shifts (Table S5).

## Discussion

We find that shifting from biomass to nutrient currencies substantially increases the proportional cost of reproduction in woody iteroparous perennials — not just because reproductive tissues require more nutrients in absolute terms, but because reproductive tissues have high nutrient concentrations combined with the fact that resorption from senescing leaves and wood shrinks the effective vegetative nutrient budget, concentrating a plant’s net annual nutrient demand in reproductive tissues. This shift is most pronounced for phosphorus, driven by the exceptional P enrichment of propagules and accessory tissues combined with high P resorption efficiency from vegetative tissues. Accessory tissues, which in this community constitute most of the reproductive biomass investment (Wenk *et al*., 2018, 2025), consume as much or more N and P as propagules themselves. Together with our prior demonstration that these species approach an RA of 100% of surplus energy as they age (Wenk *et al*., 2018; Dun *et al*., 2025), these results converge on a consistent picture: these shrubs are not modest reproducers constrained by the costs of wood production, but organisms that prioritise reproduction strongly enough that it can seriously inhibit wood and vegetative growth in mid-to-late life stages.

This matters for a longstanding debate about how much plants invest in reproduction. Ecosystem-scale estimates have consistently placed woody perennial RA below 0.2 of NPP across biomes (Malhi *et al*., 2011; Kitayama *et al*., 2015; Hanbury-Brown *et al*., 2022; Ward *et al*., 2025). Our results suggest this consensus view reflects two compounding features of ecosystem-scale accounting: RA is averaged across all age classes, diluting the high investment of mid-to-late life individuals, and the NPP denominator does not separate annual growth from standing biomass. When allocation is instead tracked across individual lifetimes and expressed as a fraction of annual growth, reproductive allocation increases. If reproductive allocation is expressed as a fraction of surplus production, it is yet higher, with these shrubs investing the vast majority of available biomass in reproduction at mid-to-late life stages (Wenk et al. 2018). These high allocation values are more consistent with measurements made on annual plants and with the expectations of eco-evolutionary theory (Iwasa & Cohen, 1989; Stearns, 1992; Wenk & Falster, 2015), rather than what the ecosystem-scale literature has implied (Bullock, 1995; Kitayama *et al*., 2015; Gougherty & Templer, 2024). Further, when investment is expressed in nutrient rather than biomass currencies, the proportional cost of reproduction increases again — both because resorption returns a substantial fraction of the nutrients invested in leaves and wood back into the annual budget, shrinking the effective vegetative nutrient demand, and because reproductive tissues, particularly propagules and floral accessory tissues, are disproportionately enriched in N and P relative to their biomass.

A second finding shapes how broadly this conclusion applies. Despite considerable variation in lifespan, height, and key functional traits across the 14 study species (Table S6) (Wenk *et al*., 2018; Dun *et al*., 2025), allocation patterns were far more consistent among species than among accounting schemes or currencies. What we conclude about reproductive investment in woody perennials depends more on how we measure it than on which species we study. This contrasts with data in a recent meta-analysis that indicated shrubs have quite variable RA, but they did not consider differences in how RA was calculated (Pérez-Martínez & Méndez, 2021).

### Tissue nutrient concentrations

Functionally similar tissues exhibited modest interspecific variation in nutrient concentration. For all vegetative and reproductive tissue categories, variation in N or P concentrations between species was far less than variation across tissues (Figure 2, Tables 1-3, Tables S2-S4). This was as expected: actively dividing tissues, such as reproductive tissues, have high mRNA concentrations and are correspondingly higher in P, while photosynthetic tissues are enriched in N (Enquist *et al*., 1999; Reich *et al*., 2009; Elser *et al*., 2010). The ranking of different plant tissues across species and growing conditions generally remains constant with regard to both their nutrient concentrations (wood lowest, then bark, leaves, and finally reproductive tissues) and nutrient ratios, following established expectations (Kerkhoff *et al*., 2006; Ågren, 2008; Elser *et al*., 2010; Minden & Kleyer, 2014; Yan *et al*., 2016; Zhao *et al*., 2020; Wang *et al*., 2022).

Some reproductive tissue types did display considerable interspecific variation, due either to variation in species nutrient allocation or our unintended bulking of dissimilar accessory tissues into a single “category”. Propagules showed the greatest interspecific variation, likely because for some species the seed could not be tidily separated from dispersal tissues, diluting the nutrient concentration. Bulked accessory tissues (Figure 2a) showed more than tenfold variation in N concentration because functionally different accessory tissues had quite different nutrient concentrations and the proportional biomass allocation to different reproductive accessory tissues differed by species. This aligns with the high variation in nutrient concentrations and N:P ratios of shrub reproductive tissues identified by Pérez-Martínez and Méndez (2021) across studies and emphasises the need to measure the nutrient concentrations on functionally different accessory tissues. Woody accessory tissues (Figures 2b, 2d) also varied widely, possibly because woody cones and woody seed pods had quite different nutrient concentrations.

Specific tissue nutrient concentrations also offered insight to unique characteristics of this community. Many species displayed a particularly high bark N concentration, suggesting either bark nutrient storage or photosynthetic bark (Cernusak & Hutley, 2011; Gong *et al*., 2024); the average bark N concentration was about 50% higher than in a recent compilation across multiple continents (Gong *et al*., 2024) supporting the results of a nearby study where 14 of 15 shrub and tree species had photosynthetic bark (Rosell *et al*., 2014). Second, Proteaceae species displayed exceptionally high seed P concentrations. P-loading of seeds in low-nutrient soils in western Australia (Lee & Fenner, 1989; Zhang *et al*., 2007; Groom & Lamont, 2010) is hypothesised to enhance germination and early establishment (Stock *et al*., 1990; Milberg & Lamont, 1997; Vaughton & Ramsey, 2001); the P contents of the soils at the study site were also low (Table S1).

N:P scaling exponents for reproductive tissues were substantially steeper than the globally established N ∝ P^2/3^ (Enquist *et al*., 1999; Kerkhoff *et al*., 2006; Reich *et al*., 2009; Elser *et al*., 2010; Zhao *et al*., 2020; Wang *et al*., 2022; Gong *et al*., 2024), driven by exceptionally high P concentrations in reproductive tissues (Figures 2a, b, Fig S1). Too few datasets exist globally to know if this steeper scaling exponent is common across many plant communities; Kerkhoff *et al*. (2006) also noted a steeper scaling factor for reproductive tissues, while Minden and Kleyer (2014) found identical scaling relationships for diaspores and leaves. For vegetative tissues, N and P were uncorrelated across taxa.

These species resorbed P from leaves, sapwood and bark (Table 3, Figure 3) at among the highest rates recorded (Brant & Chen, 2015; Freschet *et al*., 2021) — yet N resorption was unexpectedly low. Nutrient resorption offers an important mechanism for plants to internally recycle nutrients; plants are known to translocate nutrients directly from senescing vegetative material into developing propagules (Urban *et al*., 2004; Ichie & Nakagawa, 2013). In a recent study of a nearby community, Dhakal *et al*. (2025) observed that most species resorbed P to below the global threshold of 0.4 mg P/g dry mass proposed by Killingbeck (1996) for ‘complete’ P resorption in evergreen woody species; all species in this study also surpassed that threshold by high margins. However, here quite low amounts of nitrogen were resorbed (21% from leaves, on average), far lower than observed in nearby studies on similar communities, while P resorption efficiency (77% from leaves, on average) and P concentrations in senesced leaves were similar (Wright & Westoby, 2003; Dhakal *et al*., 2025) (Table 3, Figure 3). This puzzling, community-wide pattern suggests environmental conditions that made nitrogen resorption costly. Nutrient resorption from senescing leaves is known to be closely tied to soil nutrient contents and biome (McGroddy *et al*., 2004); the study sites were notably depauperate in P (1.03–2.43 mg P/g soil, Mehlich-3 extraction) but not in N (1127–3140 mg total N/g soil) (Table S4), and ostensibly retrieving N from senescing tissues was not prioritised for these species.

### Nutrient allocation accounting schemes

Accounting scheme and currency had far larger effects on perceived allocation than species identity (Figures 3, 4). While most species exhibited broadly similar allocation patterns within a single accounting scheme, the proportion of N or P allocated to each tissue shifted based on accounting scheme, due both to shifts in biomass allocation and to the relative nutrient cost of each tissue. For all accounting schemes, propagules and, to a lesser extent, reproductive accessory tissues, became relatively more costly in N and especially P currencies (Figures 4a-d, righthand panels). A snapshot harvest over-represents investment in wood and leaves as, for these evergreen perennials, much of the biomass measured at a point in time remains from a previous year’s growth (Figure 4a). Younger individuals (Figure 4b) have lower reproductive investment than do mid-aged individuals (Figure 4c), such that shifting to nutrient currencies has less pronounced effects. Accounting for nutrient resorption amplifies the relative nutrient cost of reproduction, as the P cost of vegetative tissues implicitly decreases, while plants resorb no nutrients from propagules and likely only a modest proportion from reproductive accessory tissues (Figure 4d). In contrast to the strong shift in allocation between accounting schemes and currencies, we observed a modest shift across species within a single scheme or currency.

This convergence across species has a practical implication for how measurement effort should be directed. Because allocation proportions and tissue nutrient concentrations were broadly consistent among the 14 species studied here — with between-species variation far smaller than variation attributable to accounting scheme, currency, and plant age — the dominant source of uncertainty in community-level nutrient budgets is methodological rather than biological. A study that measures all tissue types, across multiple age classes, and accounts for resorption in a small number of species will capture that methodological variation and produce more accurate community-level estimates than one that applies a snapshot harvest with bulked tissue concentrations across many species. Where resources are limited, depth of accounting matters more than breadth of species coverage.

### Belowground allocation remains unquantified

While the accounting scheme and measurements made in this study were among the most comprehensive for a community of iteroparous, perennial shrubs, they are still imperfect. The greatest shortcoming is that we could not excavate roots, which are also an important component of dry mass and nutrient budgets; the roots of these species can likely extend many metres, making excavation both impossible and inaccurate. The dearth of data on root biomass allocation traits for Australia’s flora is evident in AusTraits (Falster *et al*. 2021, v7.0.0): data on root:shoot ratio in adult, field-grown plants exist for just four shrubs, ranging from 0.13 (*Daviesia mimosoides*) to 2.54 (*Melichrus urceolatus*) (Mokany & Ash, 2008), while global average root mass fraction by biome is 0.36 for woodlands and 0.47 for shrublands. If root biomass were indeed approximately equal to aboveground biomass, reproductive allocation values would be much lower, although the shifts across ages and nutrient currencies would remain equally strong.

### Nutrient resorption only partly accounted for

We have determined that plants resorb high proportions of P and low proportions of N from sapwood and bark (Figure 3), but have not applied these resorptions to the allocation calculations due to our inability to separate sapwood from heartwood; including any nutrient resorption from woody tissues would further increase RA. We also do not have nutrient resorption values for senescing reproductive tissues; our calculations assumed a complete absence of resorption from aborted buds, flowers, and shed floral parts post-antithesis. While plants are unlikely to recover nutrients from parts shed unintentionally (herbivory, weather events), for floral parts that remain on the plant until planned senescence, the plant will likely return some nutrients. RA would decline modestly if nutrients were resorbed from senescing reproductive tissues.

### Closing remarks

Our findings suggest that for woody iteroparous perennials, reproductive tissues — especially accessory tissues — represent a large nutrient sink. Studies or models that omit or underestimate reproductive nutrient costs therefore likely overestimate the proportion of a plant’s NPP available for vegetative growth and correspondingly misrepresent a forest’s growth rate and carbon sequestration ability. That these patterns hold consistently across 14 species differing substantially in lifespan, growth form, and functional traits suggests the conclusions are not idiosyncratic to this community — the methodological artefacts depressing apparent RA in woody perennials are general, and correcting for them is likely to revise upward our understanding of reproductive investment across the biome. Together, these findings suggest that the long-standing perception of woody perennials as modest reproducers reflects the limitations of how investment has been measured rather than any fundamental difference in life history strategy — when reproduction is quantified in the currencies that matter for plant function, and across the full arc of a plant’s lifetime, these shrubs emerge as organisms that prioritise reproduction as strongly as life history theory predicts they should.

## Acknowledgements

Field and lab work was funded by the Australian Research Council via a Laureate Fellowship awarded to Mark Westoby and a Future Fellowship awarded to Daniel S. Falster. Ian J. Wright acknowledges support from the Australian Research Council (DP220102547). Fieldwork was carried out under NPWS Scientific Licence S10564, with permission from local authorities. Claude Sonnet 4.6 AI model was used for minor text editing of fully drafted paragraphs.

## Author contributions

EW, DF and MW designed the experiment. EW performed the fieldwork and lab analysis. EW and DF led the data analysis. EW led the writing. All authors contributed to manuscript revisions.

## Data availability

All raw data, code to analyse data and plot figures is available at https://github.com/traitecoevo/reproductive_allocation_kuringgai.

