## Supplementary Tables for "Nutrient currencies and P resorption greatly amplify the perceived costs of reproductive allocation in perennial heath shrubs"

Table S1. Sites sampled in Kuring’gai National Park, Sydney, NSW. Date of last fire serves as a proxy for the age of plants, as all study species are obligate seeders, germinating soon after fire.

| **Site name** | **Date of fire** | **Age at harvest (yr)** | **Longitude** | **Latitude** | **Soil C (%)** | **Soil N**  **(mg/g)** | **Soil P (mg/kg)** |
| --- | --- | --- | --- | --- | --- | --- | --- |
| Wilunga2012 | 2013 | 1.35 | 151.2622 | −33.6174 | 2.30 | 2.263 | 1.77 |
| Waratah2011 | 2011 | 2.4 | 151.2575 | −33.6408 | 1.96 | 1.597 | 2.43 |
| Basin2007 | 2007 | 5 | 151.2855 | −33.5951 | 1.36 | 1.250 | 1.2 |
| Basin2005 | 2005 | 7 | 151.2836 | −33.5933 | 1.87 | 2.207 | 1.43 |
| Waratah2003 | 2003 | 9 | 151.257 | −33.6413 | 1.73 | 1.117 | 1.03 |
| Basin1981 | 1981 | 32 | 151.2852 | −33.5954 | 2.24 | 1.127 | 1.77 |

Table S2. Vegetative tissue nutrient concentrations, by species and tissue.

a. Nitrogen (mg/g) (mean ± SE)
b. Phosphorus (mg/kg) (mean ± SE)

| **species** | **Leaves** | **Sapwood (wood from tip)** | **Wood from**  **stem base** | **Bark** |
| --- | --- | --- | --- | --- |
| *Banksia ericifolia* | 16.21 ± 1.25 | 3.66 ± 0.71 | 0.82 ± 0.03 | 3.63 ± 0.14 |
| *Boronia ledifolia* | 19.36 ± 0.40 | 2.71 ± 0.36 | 1.47 ± 0.24 | 7.05 ± 0.80 |
| *Conospermum ericifolium* | 12.48 ± 2.18 | 2.57 ± 0.49 | 3.08 ± 0.19 | 6.08 ± 0.12 |
| *Epacris microphylla* | 14.43 ± 0.86 | 2.44 ± 0.19 | 2.24 ± 0.25 | 6.86 ± 0.65 |
| *Grevillea buxifolia* | 10.75 ± 0.83 | 1.77 ± 0.31 | 0.23 ± 0.01 | 2.55 ± 0.07 |
| *Grevillea speciosa* | 11.08 ± 0.24 | 1.36 ± 0.05 | 1.56 ± 0.13 | 2.55 ± 0.32 |
| *Hakea teretifolia* | 7.40 ± 0.16 | 2.12 ± 0.31 | 2.21 ± 0.48 | 4.52 ± 0.43 |
| *Hemigenia purpurea* | 15.55 ± 0.42 | 1.78 ± 1.03 | 0.66 ± 0.32 | 5.12 ± 0.06 |
| *Leucopogon esquamatus* | 9.32 ± 0.93 | 2.99 ± 0.68 | 2.56 ± 0.18 | 5.51 ± 0.20 |
| *Persoonia lanceolata* | 9.56 ± 0.52 | 3.83 ± 0.40 | 2.29 ± 0.92 | 5.15 ± 0.17 |
| *Petrophile pulchella* | 11.31 ± 2.32 | 2.71 ± 0.09 | 1.17 ± 0.08 | 3.65 ± 0.07 |
| *Phyllota phylicoides* | 12.10 ± 2.50 | 5.36 ± 0.56 | 4.22 ± 0.06 | 10.33 ± 0.51 |
| *Pimelea linifolia* | 15.79 ± 0.39 | 2.11 ± 0.28 | 3.34 ± 0.24 | 5.64 ± 0.38 |
| *Pultenaea tuberculata* | 12.01 ± 0.89 | 5.99 ± 0.53 | 4.11 ± 0.16 | 10.96 ± 0.54 |

| **species** | **Leaves** | **Sapwood (wood from tip)** | **Wood from**  **stem base** | **Bark** |
| --- | --- | --- | --- | --- |
| *Banksia ericifolia* | 213.35 ± 37.79 | 104.56 ± 12.61 | 37.44 ± 15.84 | 85.66 ± 18.63 |
| *Boronia ledifolia* | 226.31 ± 10.60 | 82.43 ± 5.42 | 45.76 ± 7.19 | 95.67 ± 15.77 |
| *Conospermum ericifolium* | 187.15 ± 11.60 | 72.31 ± 36.77 | 57.37 ± 19.93 | 122.56 ± 46.52 |
| *Epacris microphylla* | 189.93 ± 16.19 | 77.16 ± 7.08 | 55.40 ± 8.33 | 109.52 ± 8.80 |
| *Grevillea buxifolia* | 193.57 ± 16.42 | 120.84 ± 10.77 | 80.38 ± 33.04 | 128.48 ± 27.00 |
| *Grevillea speciosa* | 209.18 ± 26.54 | 134.50 ± 43.06 | 207.33 ± 132.94 | 152.15 ± 46.49 |
| *Hakea teretifolia* | 129.34 ± 5.38 | 109.67 ± 13.86 | 58.31 ± 16.40 | 87.77 ± 13.92 |
| *Hemigenia purpurea* | 224.48 ± 49.25 | 97.54 ± 5.16 | 57.28 ± 6.90 | 106.11 ± 10.49 |
| *Leucopogon esquamatus* | 214.91 ± 27.63 | 99.63 ± 18.44 | 77.46 ± 12.93 | 104.79 ± 20.02 |
| *Persoonia lanceolata* | 247.73 ± 30.98 | 217.90 ± 53.61 | 49.51 ± 17.43 | 151.26 ± 21.14 |
| *Petrophile pulchella* | 179.77 ± 18.30 | 111.65 ± 36.22 | 65.93 ± 25.61 | 132.22 ± 37.01 |
| *Phyllota phylicoides* | 207.14 ± 22.90 | 90.64 ± 1.10 | 47.09 ± 3.22 | 102.74 ± 19.38 |
| *Pimelea linifolia* | 391.07 ± 43.86 | 70.60 ± 8.69 | 56.38 ± 11.70 | 171.77 ± 18.87 |
| *Pultenaea tuberculata* | 166.30 ± 10.74 | 151.84 ± 13.51 | 47.04 ± 2.30 | 107.48 ± 14.65 |

| **species** | **Leaves** | **Sapwood (wood from tip)** | **Wood from**  **stem base** | **Bark** |
| --- | --- | --- | --- | --- |
| *Banksia ericifolia* | 80.80 ± 7.65 | 39.92 ± 12.40 | 37.25 ± 10.44 | 51.77 ± 12.55 |
| *Boronia ledifolia* | 87.91 ± 5.02 | 37.15 ± 8.13 | 51.69 ± 21.24 | 90.07 ± 13.74 |
| *Conospermum ericifolium* | 68.65 ± 14.70 | 60.11 ± 22.42 | 79.05 ± 26.07 | 86.37 ± 33.63 |
| *Epacris microphylla* | 79.76 ± 4.91 | 35.39 ± 4.08 | 61.74 ± 22.12 | 65.85 ± 6.18 |
| *Grevillea buxifolia* | 55.80 ± 2.58 | 14.47 ± 1.88 | 4.42 ± 0.99 | 23.82 ± 4.93 |
| *Grevillea speciosa* | 55.87 ± 7.80 | 13.04 ± 3.62 | 17.50 ± 6.20 | 22.19 ± 6.18 |
| *Hakea teretifolia* | 58.58 ± 2.68 | 23.70 ± 3.63 | 75.73 ± 19.42 | 66.24 ± 10.46 |
| *Hemigenia purpurea* | 77.06 ± 12.23 | 19.10 ± 11.44 | 12.09 ± 5.91 | 49.56 ± 4.54 |
| *Leucopogon esquamatus* | 45.13 ± 4.08 | 29.42 ± 2.76 | 37.35 ± 7.04 | 61.92 ± 12.69 |
| *Persoonia lanceolata* | 41.04 ± 5.24 | 21.35 ± 4.54 | 44.61 ± 2.77 | 37.21 ± 5.74 |
| *Petrophile pulchella* | 64.94 ± 12.13 | 32.75 ± 7.08 | 29.28 ± 8.39 | 37.50 ± 9.49 |
| *Phyllota thylakoids* | 61.53 ± 14.21 | 59.36 ± 6.70 | 90.62 ± 5.14 | 109.88 ± 16.65 |
| *Pimelea linifolia* | 41.54 ± 5.34 | 31.69 ± 7.50 | 63.19 ± 11.05 | 33.51 ± 3.95 |
| *Pultenaea tuberculata* | 72.08 ± 1.32 | 41.17 ± 5.33 | 88.85 ± 7.83 | 109.77 ± 15.20 |

c. N:P ratio (mean ± SE)

Table S3. Reproductive tissue nutrient concentrations, by species and tissue.

| **species** | **accessory tissue category** | **accessory tissue** | **N (mg/g)** | **P (mg/kg)** | **NP ratio** |
| --- | --- | --- | --- | --- | --- |
| *Banksia ericifolia* | propagule | seed | 96.06 | 8205.32 | 11.71 |
| *Boronia ledifolia* | propagule | seed | 31.69 | 1387.15 | 22.85 |
| *Conospermum ericifolium* | propagule | fruit, mature | 47.99 | 6089.40 | 7.88 |
| *Epacris microphylla* | propagule | fruit, mature | 15.71 | 93.32 | 168.34 |
| *Grevillea buxifolia* | propagule | seed | 33.09 | 3725.01 | 8.88 |
| *Grevillea speciosa* | propagule | seed | 46.39 | 4190.76 | 11.07 |
| *Hakea teretifolia* | propagule | seed | 78.12 | 21455.46 | 3.64 |
| *Hemigenia purpurea* | propagule | fruit, mature | 14.50 | 1204.11 | 12.04 |
| *Leucopogon esquamatus* | propagule | fruit, mature | 50.00 | 249.11 | 200.72 |
| *Persoonia lanceolata* | propagule | seed | 5.55 | 335.73 | 16.52 |
| *Petrophile pulchella* | propagule | fruit, mature | 47.89 | 4358.22 | 10.99 |
| *Phyllota phylicoides* | propagule | seed | 50.00 | 3132.73 | 15.96 |
| *Pimelea linifolia* | propagule | seed | 50.00 | 1605.86 | 31.14 |
| *Pultenaea tuberculata* | propagule | seed | 50.05 | 3516.53 | 14.23 |
| *Banksia ericifolia* | immature propagule | fruit/seed, immature | 12.37 | 130.60 | 94.72 |
| *Boronia ledifolia* | immature propagule | fruit/seed, immature | 15.66 | 412.28 | 37.98 |
| *Epacris microphylla* | immature propagule | fruit/seed, immature | 8.89 | 393.99 | 22.56 |
| *Grevillea buxifolia* | immature propagule | fruit/seed, immature | 17.36 | 1471.53 | 11.80 |
| *Grevillea speciosa* | immature propagule | fruit/seed, immature | 14.81 | 965.18 | 15.34 |
| *Hakea teretifolia* | immature propagule | fruit/seed, immature | 49.05 | 953.22 | 51.46 |
| *Hemigenia purpurea* | immature propagule | fruit/seed, immature | 15.25 | 1546.34 | 9.86 |
| *Leucopogon esquamatus* | immature propagule | fruit/seed, immature | 10.00 | 745.36 | 13.42 |
| *Persoonia lanceolata* | immature propagule | fruit/seed, immature | 11.18 | 807.25 | 13.85 |
| *Petrophile pulchella* | immature propagule | fruit/seed, immature | 7.38 | 83.54 | 88.30 |
| *Phyllota phylicoides* | immature propagule | fruit/seed, immature | 26.48 | 1233.59 | 21.47 |
| *Pimelea linifolia* | immature propagule | fruit/seed, immature | 16.71 | 882.65 | 18.93 |
| *Banksia ericifolia* | floral accessory | bud | 16.76 | 371.97 | 45.06 |
| *Banksia ericifolia* | floral accessory | petals | 15.72 | 179.72 | 87.47 |
| *Boronia ledifolia* | floral accessory | bud | 17.12 | 630.63 | 27.15 |
| *Boronia ledifolia* | floral accessory | petals | 21.25 | 455.68 | 46.63 |
| *Conospermum ericifolium* | floral accessory | finished flower | 11.44 | 822.08 | 13.92 |
| *Conospermum ericifolium* | floral accessory | petals | 8.25 | 399.91 | 20.62 |
| *Epacris microphylla* | floral accessory | bud | 15.27 | 578.16 | 26.41 |
| *Epacris microphylla* | floral accessory | finished flower | 7.79 | 449.21 | 17.35 |
| *Epacris microphylla* | floral accessory | petals | 5.95 | 199.41 | 29.82 |
| *Grevillea buxifolia* | floral accessory | bud | 16.10 | 946.33 | 17.01 |
| *Grevillea buxifolia* | floral accessory | finished flower | 14.44 | 381.24 | 37.88 |
| *Grevillea buxifolia* | floral accessory | petals | 11.05 | 67.75 | 163.09 |
| *Grevillea speciosa* | floral accessory | bud | 20.57 | 1273.15 | 16.16 |
| *Grevillea speciosa* | floral accessory | petals | 11.01 | 200.73 | 54.85 |
| *Hakea teretifolia* | floral accessory | bud | 9.97 | 446.15 | 22.35 |
| *Hakea teretifolia* | floral accessory | flower stigma | 10.00 | 907.76 | 11.02 |
| *Hakea teretifolia* | floral accessory | petals | 17.90 | 1357.59 | 13.19 |
| *Hemigenia purpurea* | floral accessory | bud | 10.82 | 758.26 | 14.27 |
| *Hemigenia purpurea* | floral accessory | petals | 7.71 | 493.74 | 15.61 |
| *Leucopogon esquamatus* | floral accessory | bud | 10.46 | 518.67 | 20.17 |
| *Leucopogon esquamatus* | floral accessory | petals | 4.20 | 104.19 | 40.33 |
| *Persoonia lanceolata* | floral accessory | bud | 10.11 | 948.75 | 10.66 |
| *Persoonia lanceolata* | floral accessory | petals | 9.68 | 697.29 | 13.88 |
| *Petrophile pulchella* | floral accessory | petals | 6.90 | 393.44 | 17.53 |
| *Phyllota phylicoides* | floral accessory | petals | 18.11 | 587.78 | 30.81 |
| *Pimelea linifolia* | floral accessory | petals | 15.00 | 550.47 | 27.25 |
| *Pultenaea tuberculata* | floral accessory | petals | 11.52 | 303.57 | 37.95 |
| *Conospermum ericifolium* | green accessory | green parts, various | 10.88 | 297.87 | 36.53 |
| *Grevillea buxifolia* | green accessory | green parts, various | 12.06 | 903.43 | 13.35 |
| *Grevillea speciosa* | green accessory | green parts, various | 13.71 | 925.56 | 14.81 |
| *Hakea teretifolia* | green accessory | bract | 12.10 | 164.33 | 73.63 |
| *Hemigenia purpurea* | green accessory | calyx | 7.29 | 302.84 | 24.07 |
| *Leucopogon esquamatus* | green accessory | calyx | 9.37 | 371.24 | 25.25 |
| *Phyllota phylicoides* | green accessory | bract | 14.09 | 315.80 | 44.62 |
| *Phyllota phylicoides* | green accessory | calyx | 15.78 | 335.75 | 47.00 |
| *Pimelea linifolia* | green accessory | green parts, various | 12.41 | 372.29 | 33.33 |
| *Pultenaea tuberculata* | green accessory | bract | 12.68 | 82.35 | 153.98 |
| *Pultenaea tuberculata* | green accessory | calyx | 11.46 | 508.52 | 22.54 |
| *Banksia ericifolia* | woody accessory | cone | 24.43 | 234.50 | 104.18 |
| *Banksia ericifolia* | woody accessory | seed pod | 2.00 | 101.51 | 19.73 |
| *Boronia ledifolia* | woody accessory | seed pod | 13.65 | 221.10 | 61.74 |
| *Grevillea buxifolia* | woody accessory | seed pod | 5.08 | 106.89 | 47.55 |
| *Grevillea speciosa* | woody accessory | seed pod | 4.33 | 72.27 | 59.89 |
| *Hakea teretifolia* | woody accessory | seed pod | 3.69 | 146.69 | 25.17 |
| *Persoonia lanceolata* | woody accessory | seed pod | 6.47 | 514.55 | 12.57 |
| *Petrophile pulchella* | woody accessory | cone | 2.40 | 67.63 | 35.50 |
| *Phyllota phylicoides* | woody accessory | seed pod | 8.00 | 115.13 | 69.49 |
| *Pultenaea tuberculata* | woody accessory | seed pod | 6.63 | 64.72 | 102.47 |

Table S4. Senesced tissue nutrient concentrations, by species and tissue. (* samples were bulked across all individuals to obtain a sufficiently large sample size for analysis.)

a. Nitrogen (mg/g) (mean ± SE)

| **species** | **Leaves** | **Sapwood (wood from tip)** | **Bark** |
| --- | --- | --- | --- |
| *Banksia ericifolia* | 12.24 ± 0.30 | 1.43 ± 0.05 | 4.88 ± 0.08 |
| *Boronia ledifolia* | 4.51 ± 0.43 | 2.72 ± 0.29 | 5.13 ± 0.41 |
| *Conospermum ericifolium* | 9.25 ± 4.17 | 4.15* | 5.25* |
| *Epacris microphylla* | 10.94 ± 1.54 | 2.41 ± 0.25 | 5.86 ± 0.50 |
| *Grevillea buxifolia* | 8.65 ± 1.04 | 0.72 ± 0.35 | 2.28 ± 0.12 |
| *Grevillea speciosa* | 6.95 ± 0.57 | 1.27 ± 0.08 | 2.55 ± 0.20 |
| *Hakea teretifolia* | 5.40 ± 0.39 | 1.80 ± 0.21 | 3.30 ± 0.35 |
| *Hemigenia purpurea* | 16.61 ± 0.50 | 1.47 ± 0.45 | 4.78 ± 0.05 |
| *Leucopogon esquamatus* | 7.51 ± 1.73 | 2.67 ± 0.12 | 5.84 ± 0.38 |
| *Persoonia lanceolata* | 11.60 ± 1.12 | 2.66 ± 0.08 | 5.62 ± 0.10 |
| *Petrophile pulchella* | 11.56 ± 1.70 | 2.47* | 4.13* |
| *Phyllota phylicoides* | 14.27 ± 1.79 | 5.38 ± 0.24 | 13.41 ± 0.52 |
| *Pimelea linifolia* | 6.39 ± 0.38 | 4.12* | 4.00* |
| *Pultenaea tuberculata* | 8.64 ± 0.00 | 4.30 ± 0.34 | 13.13 ± 0.50 |

b. Phosphorus (mg/kg) (mean ± SE)

| **species** | **Leaves** | **Sapwood (wood from tip)** | **Bark** |
| --- | --- | --- | --- |
| *Banksia ericifolia* | 17.08 ± 1.48 | 2.16 ± 0.27 | 22.95 ± 6.54 |
| *Boronia ledifolia* | 54.17 ± 8.25 | 7.64 ± 1.38 | 52.45 ± 5.66 |
| *Conospermum ericifolium* | 69.32 ± 21.41 | 15.93* | 65.08* |
| *Epacris microphylla* | 61.38 ± 25.99 | 28.78 ± 4.94 | 70.18 ± 8.96 |
| *Grevillea buxifolia* | 76.61 ± 37.61 | 21.99 ± 1.27 | 71.91 ± 12.35 |
| *Grevillea speciosa* | 26.82 ± 6.05 | 24.78 ± 5.24 | 59.48 ± 3.56 |
| *Hakea teretifolia* | 21.03 ± 2.57 | 7.77 ± 2.27 | 30.76 ± 6.77 |
| *Hemigenia purpurea* | 53.53 ± 4.94 | 18.19 ± 6.62 | 52.83 ± 13.56 |
| *Leucopogon esquamatus* | 40.07 ± 17.28 | 42.98 ± 6.44 | 91.58 ± 19.99 |
| *Persoonia lanceolata* | 53.39 ± 13.46 | 13.38 ± 1.17 | 42.06 ± 2.84 |
| *Petrophile pulchella* | 22.19 ± 0.92 | 24.56* | 60.40* |
| *Phyllota phylicoides* | 43.75 ± 9.41 | 27.03 ± 12.15 | 53.87 ± 18.56 |
| *Pimelea linifolia* | 159.27 ± 43.92 | 26.41* | 92.37* |
| *Pultenaea tuberculata* | 31.45 ± 4.32 | 29.35 ± 9.83 | 67.06 ± 13.97 |

c. N:P ratio (mean ± SE)

| **species** | **Leaves** | **Sapwood (wood from tip)** | **Bark** |
| --- | --- | --- | --- |
| *Banksia ericifolia* | 720.13 ± 44.37 | 251.26 ± 71.23 | 688.29 ± 113.74 |
| *Boronia ledifolia* | 86.37 ± 21.01 | 109.75 ± 18.75 | 546.97 ± 173.43 |
| *Conospermum ericifolium* | 167.94 ± 111.96 | 80.63* | 260.41* |
| *Epacris microphylla* | 204.26 ± 61.43 | 89.37 ± 11.24 | 96.61 ± 16.68 |
| *Grevillea buxifolia* | 140.07 ± 55.21 | 35.33 ± 7.64 | 33.43 ± 16.50 |
| *Grevillea speciosa* | 268.15 ± 39.24 | 42.85 ± 1.11 | 57.56 ± 15.23 |
| *Hakea teretifolia* | 263.24 ± 50.78 | 161.16 ± 42.60 | 345.62 ± 59.74 |
| *Hemigenia purpurea* | 313.70 ± 38.18 | 96.53 ± 23.83 | 103.72 ± 62.40 |
| *Leucopogon esquamatus* | 253.08 ± 152.39 | 72.96 ± 21.15 | 65.73 ± 12.02 |
| *Persoonia lanceolata* | 237.68 ± 80.92 | 134.10 ± 6.69 | 200.83 ± 23.26 |
| *Petrophile pulchella* | 524.79 ± 98.56 | 68.29* | 100.55* |
| *Phyllota phylicoides* | 332.74 ± 30.65 | 315.74 ± 106.34 | 282.40 ± 93.37 |
| *Pimelea linifolia* | 42.72 ± 9.38 | 43.30* | 155.83* |
| *Pultenaea tuberculata* | 279.98 ± 38.48 | 222.21 ± 46.53 | 196.32 ± 58.77 |

Table S5. Reproductive allocation, considering different calculations and currencies.

| **species** | **RA, dry mass basis** | **RA,**  **N currency** | **RA,**  **P currency** |
| --- | --- | --- | --- |
| *Banksia ericifolia* | 0.029 | 0.068 | 0.105 |
| *Boronia ledifolia* | 0.213 | 0.353 | 0.532 |
| *Conospermum ericifolium* | 0.037 | 0.104 | 0.339 |
| *Epacris microphylla* | 0.101 | 0.205 | 0.241 |
| *Grevillea buxifolia* | 0.040 | 0.103 | 0.145 |
| *Grevillea speciosa* | 0.026 | 0.065 | 0.119 |
| *Hakea teretifolia* | 0.035 | 0.110 | 0.194 |
| *Hemigenia purpurea* | 0.074 | 0.096 | 0.238 |
| *Leucopogon esquamatus* | 0.126 | 0.357 | 0.311 |
| *Persoonia lanceolata* | 0.032 | 0.043 | 0.082 |
| *Petrophile pulchella* | 0.012 | 0.030 | 0.084 |
| *Phyllota phylicoides* | 0.042 | 0.091 | 0.158 |
| *Pimelea linifolia* | 0.076 | 0.193 | 0.251 |
| *Pultenaea tuberculata* | 0.053 | 0.080 | 0.113 |

a. Snapshot harvest (depicted in Figure 4a)
b. Vegetative investment considering only yearly growth, measured on a plant during early reproductive years (depicted in Figure 4b)

| **species** | **RA, dry mass basis** | **RA,**  **N currency** | **RA,**  **P currency** |
| --- | --- | --- | --- |
| *Banksia ericifolia* | 0.029 | 0.068 | 0.105 |
| *Boronia ledifolia* | 0.213 | 0.353 | 0.532 |
| *Conospermum ericifolium* | 0.037 | 0.104 | 0.339 |
| *Epacris microphylla* | 0.101 | 0.205 | 0.241 |
| *Grevillea buxifolia* | 0.040 | 0.103 | 0.145 |
| *Grevillea speciosa* | 0.026 | 0.065 | 0.119 |
| *Hakea teretifolia* | 0.035 | 0.110 | 0.194 |
| *Hemigenia purpurea* | 0.074 | 0.096 | 0.238 |
| *Leucopogon esquamatus* | 0.126 | 0.357 | 0.311 |
| *Persoonia lanceolata* | 0.032 | 0.043 | 0.082 |
| *Petrophile pulchella* | 0.012 | 0.030 | 0.084 |
| *Phyllota phylicoides* | 0.042 | 0.091 | 0.158 |
| *Pimelea linifolia* | 0.076 | 0.193 | 0.251 |
| *Pultenaea tuberculata* | 0.053 | 0.080 | 0.113 |

c. Vegetative investment considering only yearly growth, measured on a plant during peak reproductive years (depicted in Figure 4c)
d. Vegetative investment considering only yearly growth, measured on a plant during peak reproductive years, with nutrient resorption from leaves taken into account (depicted in Figure 4d)

| **species** | **RA, dry mass basis** | **RA,**  **N currency** | **RA,**  **P currency** |
| --- | --- | --- | --- |
| *Banksia ericifolia* | 0.085 | 0.225 | 0.883 |
| *Boronia ledifolia* | 0.555 | 0.883 | 0.973 |
| *Conospermum ericifolium* | 0.135 | 0.353 | 0.869 |
| *Epacris microphylla* | 0.418 | 0.587 | 0.827 |
| *Grevillea buxifolia* | 0.506 | 0.702 | 0.910 |
| *Grevillea speciosa* | 0.102 | 0.280 | 0.785 |
| *Hakea teretifolia* | 0.123 | 0.345 | 0.889 |
| *Hemigenia purpurea* | 0.173 | 0.218 | 0.789 |
| *Leucopogon esquamatus* | 0.299 | 0.636 | 0.810 |
| *Persoonia lanceolata* | 0.207 | 0.267 | 0.834 |
| *Petrophile pulchella* | 0.033 | 0.046 | 0.457 |
| *Phyllota phylicoides* | 0.161 | 0.291 | 0.764 |
| *Pimelea linifolia* | 0.238 | 0.601 | 0.740 |
| *Pultenaea tuberculata* | 0.123 | 0.232 | 0.651 |

| **species** | **RA, dry mass basis** | **RA,**  **N currency** | **RA,**  **P currency** |
| --- | --- | --- | --- |
| *Banksia ericifolia* | 0.085 | 0.162 | 0.296 |
| *Boronia ledifolia* | 0.555 | 0.774 | 0.855 |
| *Conospermum ericifolium* | 0.135 | 0.315 | 0.656 |
| *Epacris microphylla* | 0.418 | 0.534 | 0.621 |
| *Grevillea buxifolia* | 0.506 | 0.646 | 0.775 |
| *Grevillea speciosa* | 0.102 | 0.203 | 0.351 |
| *Hakea teretifolia* | 0.123 | 0.286 | 0.483 |
| *Hemigenia purpurea* | 0.173 | 0.213 | 0.447 |
| *Leucopogon esquamatus* | 0.299 | 0.591 | 0.534 |
| *Persoonia lanceolata* | 0.207 | 0.243 | 0.385 |
| *Petrophile pulchella* | 0.033 | 0.045 | 0.119 |
| *Phyllota phylicoides* | 0.161 | 0.291 | 0.440 |
| *Pimelea linifolia* | 0.238 | 0.441 | 0.531 |
| *Pultenaea tuberculata* | 0.123 | 0.179 | 0.263 |

Table S6. Functional trait values for each species across study sites, described in greater detail in Wenk 2018, Dun 2025. The propagule includes the mass of the enmbryo and endosperm.

| **species** | **Propagule mass**  **(mg)** | **Leaf mass per area (g/m2)** | **Wood**  **Density (g/cm3)** | **Maximum height**  **(m)** |
| --- | --- | --- | --- | --- |
| *Banksia ericifolia* | 24.06 | 224 | 0.59 | 2.79 |
| *Boronia ledifolia* | 2.10 | 159 | 0.86 | 0.77 |
| *Conospermum ericifolium* | 0.69 | 206 | 0.79 | 0.99 |
| *Epacris microphylla* | 0.02 | 122 | 0.73 | 1.37 |
| *Grevillea buxifolia* | 26.70 | 146 | 0.73 | 1.37 |
| *Grevillea speciosa* | 13.48 | 169 | 0.74 | 1.00 |
| *Hakea teretifolia* | 8.18 | 516 | 0.57 | 3.01 |
| *Hemigenia purpurea* | 0.30 | 205 | 0.83 | 0.71 |
| *Leucopogon esquamatus* | 0.81 | 129 | 0.79 | 0.99 |
| *Persoonia lanceolata* | 14.39 | 203 | 0.67 | 2.14 |
| *Petrophile pulchella* | 2.21 | 297 | 0.66 | 2.01 |
| *Phyllota phylicoides* | 1.71 | 174 | 0.85 | 1.63 |
| *Pimelea linifolia* | 24.06 | 224 | 0.59 | 2.79 |
| *Pultenaea tuberculata* | 2.10 | 159 | 0.86 | 0.77 |
